# A high content imaging assay for identification of specific inhibitors of native *Plasmodium* liver stage protein synthesis

**DOI:** 10.1101/2024.05.29.596519

**Authors:** James L. McLellan, Beatriz Morales-Hernandez, Sarah Saeger, Kirsten K. Hanson

## Abstract

*Plasmodium* parasite resistance to antimalarial drugs is a serious threat to public health in malaria-endemic areas. Compounds that target core cellular processes like translation are highly desirable, as they should be multistage actives, capable of killing parasites in the liver and blood, regardless of molecular target or mechanism. Assays that can identify these compounds are thus needed. Recently, specific quantification of native *Plasmodium berghei* liver stage protein synthesis as well as that of the hepatoma cells supporting parasite growth, was achieved via automated confocal feedback microscopy of the o-propargyl puromycin (OPP)-labeled nascent proteome, but this imaging modality is limited in throughput. Here, we developed and validated a miniaturized high content imaging (HCI) version of the OPP assay that increases throughput, before deploying this approach to screen the Pathogen Box. We identified only two hits, both of which are parasite-specific quinoline-4-carboxamides, and analogues of the clinical candidate and known inhibitor of blood and liver stage protein synthesis, DDD107498/cabamiquine. We further show that these compounds have strikingly distinct relationships between their antiplasmodial and translation inhibition efficacies. These results demonstrate the utility and reliability of the *P. berghei* liver stage OPP HCI assay for specific, single-well quantification of *Plasmodium* and human protein synthesis in the native cellular context, allowing identification of selective *Plasmodium* translation inhibitors with the highest potential for multistage activity.

## Introduction

*Plasmodium* parasites have a two-host, multistage lifecycle, with human infection beginning when sporozoites are transferred via anopheline mosquito bite. These sporozoites migrate in the skin until entering the bloodstream, and are swept to the liver where they arrest, exit the circulation, and actively invade a hepatocyte inside a vacuole derived from the hepatocyte’s own plasma membrane. Once inside this parasitophorous vacuole, the parasite will undergo a dramatic period of growth and replication, before forming thousands of hepatic merozoites(1). The liver stage ends with the release of these hepatic merozoites back into the bloodstream, once again wrapped in host cell plasma membrane as merosomes, where they will eventually invade erythrocytes, commencing the blood stage (2-4). Unlike the single-pass liver stage, which functions as an amplification step in infection, most blood stage parasites engage in cyclical asexual replication, and it is this continuous process of erythrocyte invasion, parasite replication, and parasite egress via erythrocyte rupture, which causes the disease malaria (5). A minority of blood stage parasites are developmentally committed to sexual replication, and these gametocytes remain nonreplicative until they are taken up via mosquito bite and undergo sexual replication while inside the bloodmeal in the mosquito midgut (6). Antimalarial drugs were first developed for chemotherapy, and thus must target the asexual blood stage. While chemotherapy remains crucial, current antimalarial drug discovery aims to maximize the utility of new drugs by identifying compounds that also target the other human stages of the parasite (7, 8). A compound that targets the liver stage is arguably ideal for antimalarial prophylaxis, as killing the parasites before they reach the blood not only protects the infected individual from disease, but also prevents that individual from transmitting the parasite back to mosquitos, thus enhancing protection at the population level.

High throughput cellular screens have identified many compounds capable of killing both the asexual blood stage (ABS) and the liver stage (LS) parasites (9), and amongst the promising compound classes, as defined by cellular mechanism, are those that inhibit cytoplasmic *Plasmodium* protein synthesis. In addition to the nuclear genome, *Plasmodium* parasites have endosymbiont-derived genomes, with that of the apicoplast encoding genes for an essentially bacterial translational machinery, which is used to synthesize proteins from the mRNAs transcribed from the apicoplast genome, and is successfully targeted by a range of antibacterials drugs (10, 11). The cytoplasmic translation apparatus, which produces proteins from mRNAs of nucleus-encoded genes, is eukaryotic in nature, and responsible for the overwhelming bulk of all *Plasmodium* translation (12, 13). The majority of the parasite-specific cytoplasmic translation inhibitors identified to date act by inhibiting amino-acyl tRNA synthetase (aaRS) enzymes that aminoacylate tRNAs in the cytoplasm (14, 15), while the DDD107498 (also known as M5715, and in currently in clinical development as cabamiquine targets *Plasmodium* eEF2, a highly conserved elongation factor required to catalyze the translation elongation process, based on evolved resistance *in vitro*, in mice, and in human subjects (16-18). Given the promise of translation inhibitors as multistage active antimalarials, approaches to identifying them have taken advantage of both enzymatic target-based screens (14, 19) and *P. falciparum* lysate-based *in vitro* translation assays screens (20-22). LS parasites reside in highly translationally active hepatocytes and no protocol exists for their isolation, thus direct assessment of LS translation and its pharmacological inhibition was only recently achieved with a quantitative bioimaging approach (23). In this method, *P. berghei*-infected HepG2 cell monolayers, a mainstay LS drug discovery infection model, are exposed to a modified puromycin analogue, o-propargyl puromycin (OPP) (24), a charged tRNA mimetic, that is incorporated into both human and parasite nascent and terminates them; subsequent click chemistry attaches a fluorophore to each OPP molecule, thus allowing visualization and quantification of the HepG2 and *P. berghei* nascent proteomes. The OPP-labeled nascent proteomes are images via automated confocal feedback microscopy (ACFM)(25), which allows acquisition of large, unbiased datasets of single parasite images for downstream computational separation of host and parasite signals, and the quantification of each. ACFM is time consuming, though, and a higher throughput approach is needed to support efficient compound screening to identify translation inhibitors. Here, we validate a novel pseudo-widefield, 384-well plate (384wp) high content imaging (HCI) approach using samples imaged with both HCI and ACFM modalities. We then run pilot screens of the Medicines for Malaria Venture (MMV) validation set and the Pathogen Box to demonstrate the promise of this approach for efficient medium throughput identification of *Plasmodium* liver stage translation inhibitors, identifying two Pathogen Box compounds as *Plasmodium* translation inhibitors in the native cellular context.

## Results and Discussion

### Development and validation of the OPP HCI assay

We set out to develop a simplified OPP HCI assay, capable of providing a reliable readout of *Plasmodium* liver stage translational output and allowing identification of known and novel translation inhibitors that could be implemented on a wide variety of microscopes or imaging systems. To this end, we used a 10X air objective to acquire pseudo-widefield images of OPP-labelled *P. berghei*-infected HepG2 monolayers of many of the compound-treated samples that we used to establish our ACFM approach (23). An image acquisition matrix was used to acquire four non-overlapping images of infected monolayers, mimicking the image acquisition layout that would be used each well of a 384wp, our desired eventual screening format. Though *P. berghei* LS parasites, also known as exoerythrocytic forms (EEFs), are merely small dots on visual inspection, 1024x1024 pixel images provided sufficient resolution to accurately segment 28 hours post-infection (hpi) parasites (Fig. 1), due to the high specificity of the anti-PbHSP70 antibody used for EEF identification. HepG2 nuclei could also be reliably segmented based on Hoechst labeling for downstream use in quantification of the *H. sapiens* nascent proteome, as previously validated (23). As in ACFM images, many of the brightest pixels in the OPP-Alexafluor555 (OPP-A555) labelled nascent proteome HCI images are from the EEFs in control wells (Fig. 1A). Treatment with anisomycin, a pan-eukaryotic active elongation inhibitor, can be seen to inhibit both *H. sapiens* and *P. berghei* translation (Fig. 1B), while treatment with *Plasmodium*-specific DDD107498 only inhibits *P. berghei* translation (Fig. 1C).

**Figure 1.**
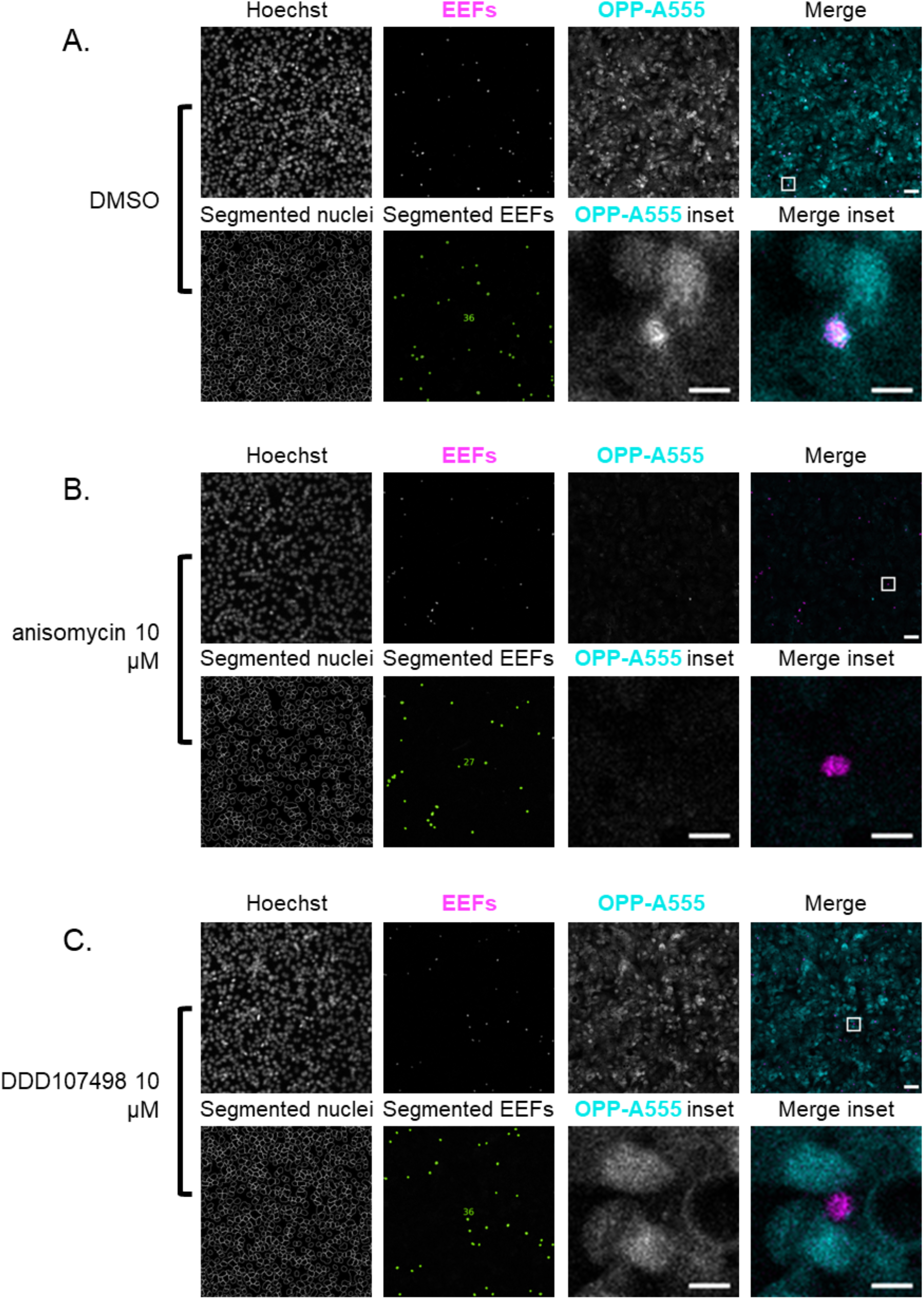
Visualization of the *Plasmodium berghei* liver stage parasite nascent proteome with high content imaging. Representative image sets of *P. berghei-*infected HepG2 at 28 hpi following a 30-minute competitive OPP labeling (coOPP) with DMSO vehicle control (**A**), and positive controls anisomycin (**B**) and DDD107498 (**C**). (**A-C**)Single channel images in greyscale show Hoechst labeled DNA, exoerythrocytic forms (EEFs) marked by anti-PbHSP70, and the nascent proteome visualized by OPP-A555 labeling. Pseudocolored merges show EEFs in magenta and OPP-A555 in cyan. Images of segmented HepG2 nuclei and segmented EEFs were generated in CellProfiler with the number of EEF objects identified reported in-image. Merged image scale bars = 50 μm for full size images (top row), and 10 μm for cropped insets.

The key challenge inherent in using this type of image for quantification is that there will be unavoidable HepG2-derived signal confounding specific quantification of the parasite nascent proteome. We reasoned that we might be able to minimize the contribution of the signal from the host nascent proteome by more aggressive shrinking of the EEF objects identified during image segmentation than we utilize for ACFM images (23, 26). Thus, we developed a segmentation and data analysis workflow to quantify parasite translation in pseudo-widefield images applying seven different computational shrinking strategies to each EEF object (Fig. 2A). We then compared our normalized average OPP-A555 mean fluorescence intensity (MFI) derived from each of the seven EEF shrinks to the matched ACFM OPP-A55 MFI for 150 individual wells, comprising DMSO controls and both active and inactive compounds at a range of concentrations. Analyzing the correlation between HCI and ACFM measurements of parasite translational intensity, we found that the most drastic shrink, in which each EEF object is reduced to only its central pixel, produced the best correlation to the ACFM data (Fig. 2A), so we chose this approach for the OPP HCI assay, and used it for all subsequent experiments. Looking at the single pixel HCI vs. ACFM data for *P. berghei* liver stage translation in a scatterplot (Fig. 2B), it can be observed that inactive treatments and controls tend to have random variation, while active treatments have lower translational intensities in the ACFM images than the HCI images. A similar trend is seen when comparing HepG2 translation intensities between the HCI and ACFM datasets (Fig. S1A), likely reflecting higher background, as expected. We extracted the concentration-response experiments for previously identified active compounds (23) from the complete dataset, and were able to detect concentration-dependent inhibition of parasite translation for our previously identified indirect and direct inhibitors of *P. berghei* translation (Fig. 2C) as well as HepG2 translation (Fig. S1B); despite the data sparsity, with only 5 concentrations tested in these experiments, calculated EC_50s_ were also generally well-correlated between HCI and ACFM datasets for both *Plasmodium* liver stage (Fig. 2D and Table S1) and HepG2 (Fig. S1C and Table S1) translation inhibition.

**Figure 2.**
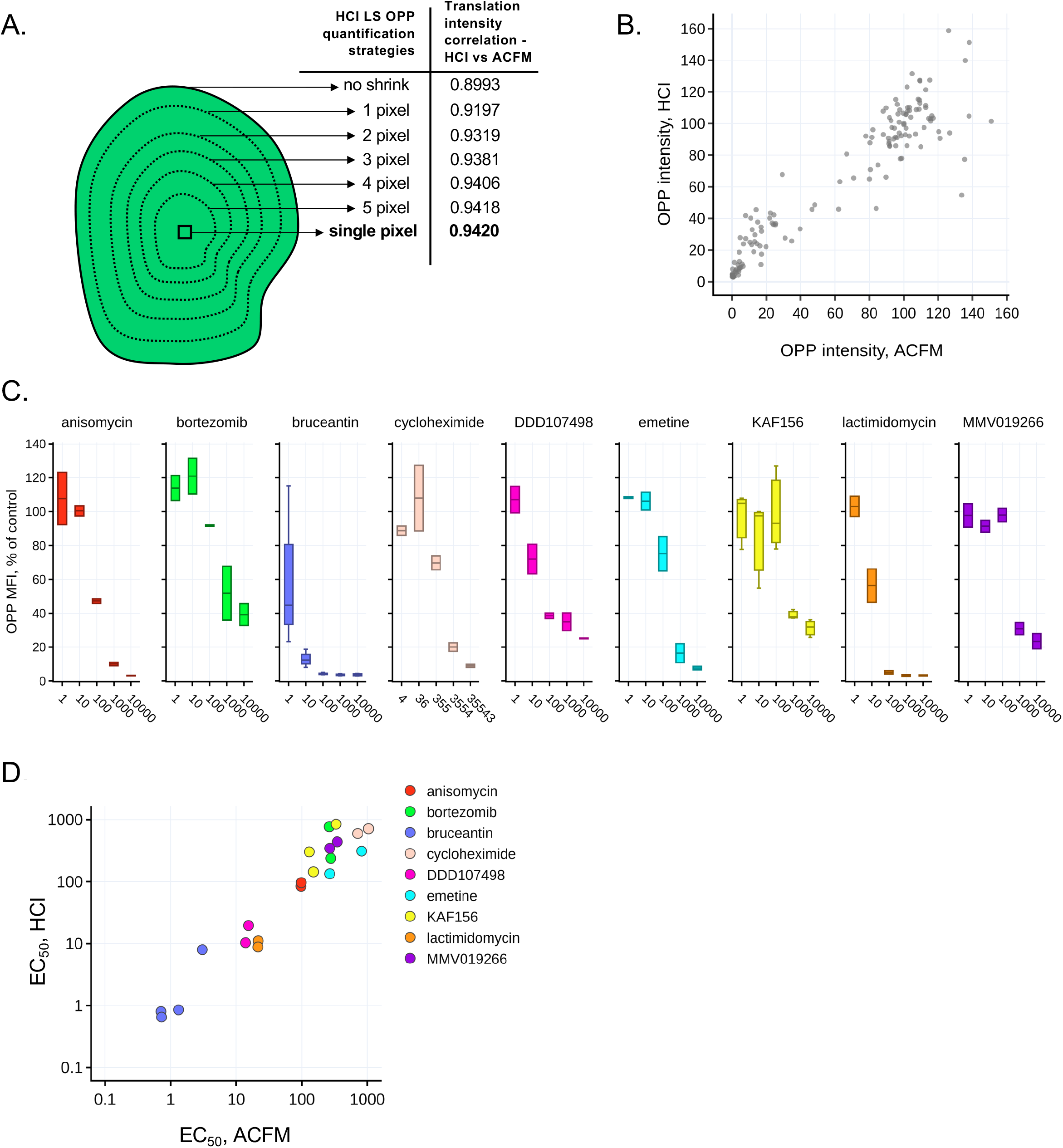
Validation of OPP HCI assay by comparison to OPP ACFM data. **A)**EEF object shrink strategies tested for correlation of EEF translation intensity (OPP-A555) in HCI images compared with published EEF translation intensities from ACFM images of the same wells (McLellan et al., 2023). **B)** Four hour (h) acute pre-treatment complete dataset, comprising 150 wells containing a range of active and inactive compounds and controls from 8 independent experiments, with single pixel HCI OPP MFI vs. the ACFM OPP MFI; each dot represents the mean OPP-A555 intensity for a single well normalized to the mean of the in-plate DMSO controls, which was set to 100. **C)** Boxplots for translation inhibition concentration-response experiments extracted from the full dataset showing HCI assay OPP MFI, normalized to in-plate DMSO controls. **D)** Comparison of translation inhibitor potency (EC_50_ value) between HCI assay and ACFM assay; plot is shown with log-log axes. For C-D), data represent 2-4 independent experiments.

Given the reliable performance of the OPP HCI assay on this dataset, we moved on to screening biologically focused compound sets for *P. berghei* liver stage translation inhibition in 384wp format. We first ran the Medicines for Malaria Venture (MMV) Validation set, a collection of 96 well-characterized antiplasmodial compounds with both structural and mechanistic diversity, at 5 μM in a competitive OPP (coOPP) assay format in which OPP is added together with the screening compounds for 30 minutes (23). The assay was run in two replicate plates, with anisomycin and DDD107498 as active controls, which inhibited parasite translation by ∼90% and ∼72% respectively (Fig. 3A). The mean Z’ (0.64 – DDD107498, 0.78 – anisomycin) and SSMD (11.1 – DDD107498, 17.4 – anisomycin) strongly supporting assay reliability. Three compounds from the library were detected as inhibitors of *P. berghei* liver stage protein synthesis, and unblinding revealed that these were the only three known translation inhibitors in the MMV validation set: DDD107498, cycloheximide, and cladosporin (Fig. 3A). DDD107498 from the library showed identical efficacy to our DDD107498 controls, and cladosporin translation inhibition efficacy was similar to that of DDD107498, while cycloheximide showed slightly greater efficacy. The OPP HCI assay clearly identifies cycloheximide as inhibiting both human and *Plasmodium* translation, like anisomycin, while DDD107498 and cladosporin are both parasite-specific (Fig. 3B). Cycloheximide is of course a well-known pan-eukaryotic translation elongation inhibitor (27, 28) and was included in the our initial validation set ((23) and Fig. 2 C,D and Fig. S1 B-C). Cladosporin, a fungal secondary metabolite, was identified in phenotypic screens as having antiplasmodial activity against both *P. falciparum* asexual blood stages (ABS) and *P. yoelii* liver stages (29, 30), and was found to inhibit cytoplasmic protein synthesis in *P. falciparum* ABS via inhibition of the cytoplasmic lysyl tRNA synthetase (cKRS) (31). Specific inhibition of *P. berghei* liver stage translation by 5 μM cladosporin was confirmed by confocal microscopy (Fig. 3C).

**Figure 3.**
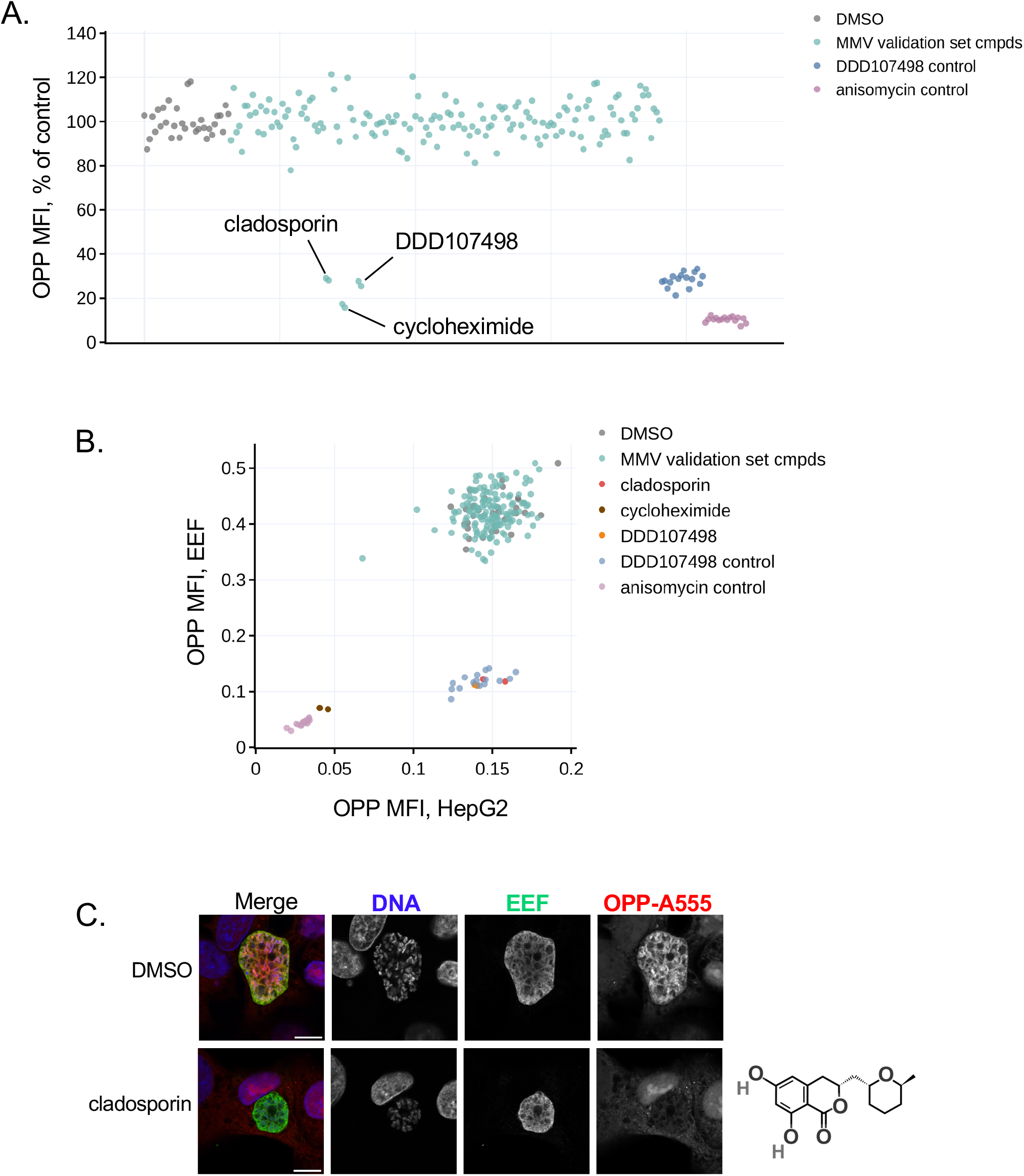
Screening the MMV Validation set. **A)** *P. berghei* translation inhibition (OPP MFI), normalized to DMSO controls following coOPP treatment with MMV validation compound set tested at 5 μM, and positive DDD107498 and anisomycin controls, tested at 1 μM and 10 μM respectively. Each dot represents a single well from the screen run in duplicate plates, and each of 3 hits are annotated in plot. All data is normalized to the mean of the in-plate DMSO controls. **B)** Mean translation intensity in EEFs compared to translation intensity from same-well HepG2 cells. **C)** Representative confocal images of the labeled *P. berghei* nascent proteome (OPP-A555) following a coOPP treatment with cladosporin (5μM) and DMSO at 47.5-48 hpi. Scale = 5μm.

### Screening the Pathogen Box and hit confirmation

We next screened the Pathogen Box, a 400-compound library comprising diverse compounds with validated activity against a range of pathogens causing neglected tropical diseases (32, 33). We again screened in technical duplicate plates at 5 μM concentration with anisomycin and DDD107498 as controls. Only two compounds from the library inhibited *P. berghei* protein synthesis: MMV634140 and MMV667494 (Fig. 4A and Table S2). Both hits had *Plasmodium* translation inhibition efficacy similar to that of DDD107498, and both were highly parasite-specific (Fig. 4A-B). Again, the assay performance was quite reliable, with mean Z’ of 0.64 and 0.45 and SSMD of 11.8 and 8.0 for anisomycin and DDD107498 respectively.

**Figure 4.**
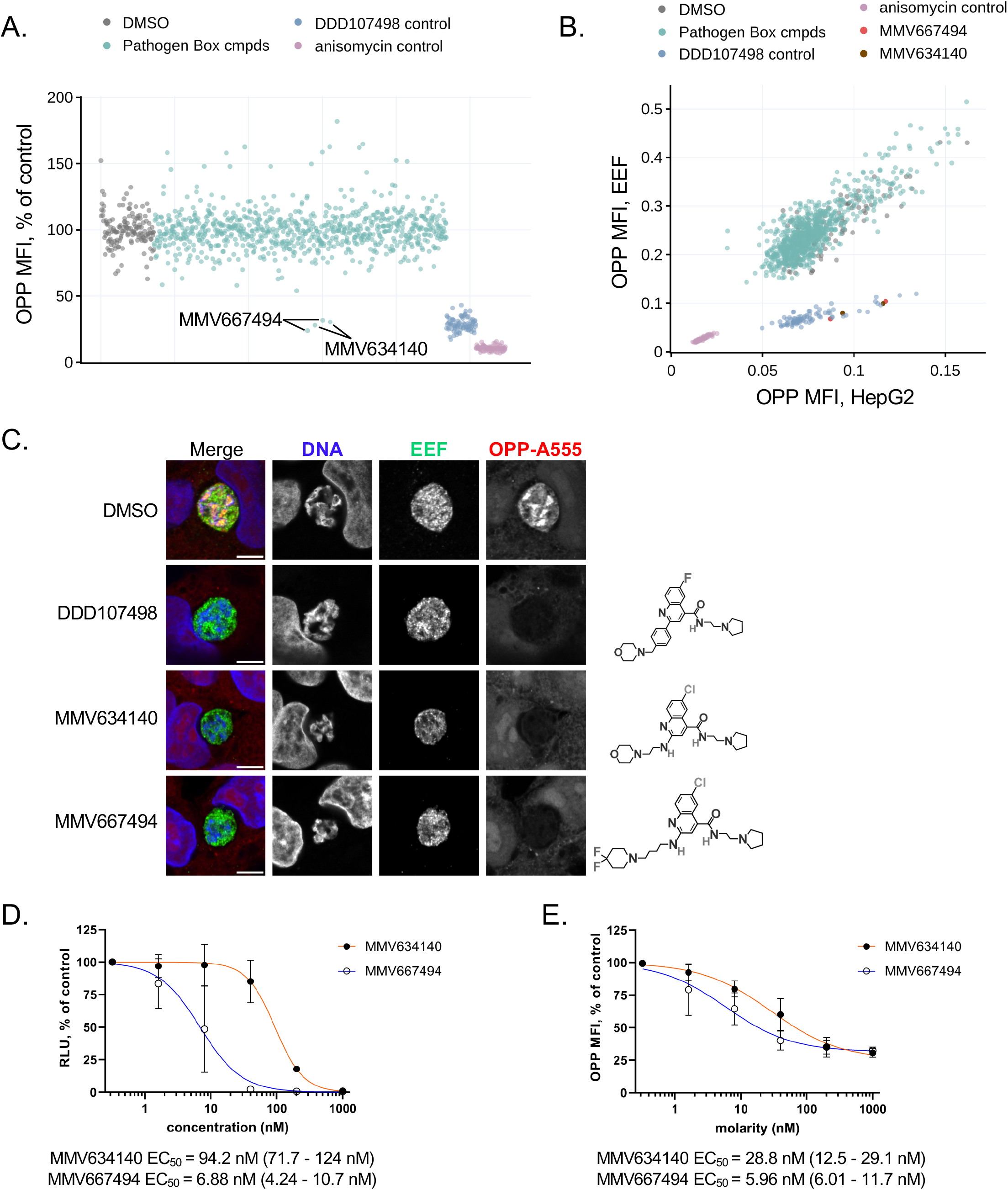
Screening the Pathogen Box. **A)** *P. berghei* liver stage translation (OPP MFI), normalized to DMSO controls following coOPP treatment with Pathogen Box library tested at 5 μM, and positive DDD107498 and anisomycin controls, at 1 μM and 10 μM respectively. Each dot represents OPP MFI from a single well, with the library screened in duplicate plates; hit compounds are annotated in plot. **B)** Mean translation intensity (OPP MFI) in EEFs compared to same-well HepG2 cells. **C)** Representative, confocal images of the *P. berghei* nascent proteome (OPP-A555) following a coOPP treatment with DMSO control, DDD107498, MMV634140 and MMV667494 (all at 10 μM) at 28 hpi. Scale bar = 5μm. **D)** Liver stage antiplasmodial activity concentrationresponse, with effects of treatment from 2-48hpi quanitfied by live luciferase assay, and **E)** Liver stage translation inhibition concentration-response using a coOPP treatment from 27.5-28 hpi. For A) and D-E), all data were normalized to the mean of the in-plate DMSO controls. In D-E), points represent the mean of three independent experiments and error bars show standard deviations. EC_50_ values and 95% confidence intervals (in parentheses) are reported for each compound.

The two hits, MMV634140 and MMV667494, are both quinoline-4-carboxamides that are structural analogues of DDD107498 (34), and are both specific inhibitors of *P. berghei* LS translation (Fig. 4C). We assessed the *P. berghei* liver stage antiplasmodial potency of both hits in 5-point, 3-fold concentration response, with the maximum concentration tested chosen based on the *P. falciparum* asexual blood stage EC_50_s provided by MMV for the Pathogen Box (33). MMV667494 had the more potent LS antiplasmodial activity of the two, with an EC_50_ of 6.8 nM, compared with 94.2 nM for MMV634140 (Fig. 4D), an approximately 14-fold difference in potency. We similarly determined the translation inhibition potency of the two hit compounds and found that their potencies were markedly more similar in this assay (Fig. 4E). MMV667494 was again the more potent compound, with an EC_50_ of 5.96 nM, while MMV634140 had a 28.8 nm EC_50_, an approximately 4.8-fold difference in liver stage translation inhibition potency (Fig. 4E). MMV634140 and MMV667494 showed essentially identical translation inhibition efficacy of ∼70% at 1 μM in the OPP HCI (Fig. 4E), just as they did at the 5 μM screening concentration (Fig. 4A-B). Translation inhibition efficacy data from the OPP HCI assay is useful in terms of relative comparisons between compounds, but due to the nature of the image acquisition used, does not provide the best readout of compound efficacy at inhibiting protein synthesis, as all parasites are not individually focused and HepG2 nascent proteome signal of course confounds measurements. We used ACFM imaging of single parasites to assess the efficacy of MMV634140 and MMV667494, and found that they have effectively identical translation inhibition efficacies of 89.7% and 86% respectively (Fig. S2). These are very similar to the efficacy of DDD107498 at 5x-and 10x its translation inhibition EC_50_ (35). The modest difference in translation inhibition potency between the two compounds seen in the OPP HCI assay was confirmed by quantification of single parasite ACFM images (Fig. S2), further validating the performance of the OPP HCI assay.

Our recent study interrogating LS parasite responses to a four-hour, acute treatment with DDD107498 and four other translation inhibitors at equivalent effective concentrations provided clear data that a compound’s efficacy in inhibiting LS translation during the treatment period does not correlate with the antiplasmodial effects that treatment causes after compound washout (35). DDD107498 was the least efficacious translation inhibitor at both at 5x- and 10x translation inhibition EC_50_, but the most efficacious antiplasmodial of the five tested, based on analysis of three days’ worth of merosome/detached cell release. Further, DDD107498 treatment of parasites that were already translationally arrested prior to compound addition exerted a marked increase in the antiplasmodial effect of the translation inhibitors alone (35). For parasites where protein synthesis was arrested with anisomycin, LysRS-IN2, or bruceantin for one hour, followed by addition of 20 nM DDD107498, then rigorous compound washout three hours later, the marked cytostatic effect of DDD107498 alone on merosome release was imposed on all the co-treated parasites. Additionally, LS developmental success (total merosome/detached cell production out to 120 hpi) was reduced by 50% for all four mechanistically diverse protein synthesis inhibitors tested, including MMV019266, upon addition of 20 nM DDD107498 to the translationally arrested parasites.

In the context of these results, the disconnect between antiplasmodial potency and translation inhibition potency seen here for MMV634140 is quite striking. The compound is approximately 3-fold more potent an inhibitor of LS protein synthesis than it is of LS growth (Figs 4D-E).DDD107498 also has clear a disconnect between its antiplasmodial potency and translation inhibition potency that spans *Plasmodium* stages and species, though it has the opposite directionality from that of MMV634140. *P. falciparum* asexual blood stage (ABS) translation has been quantified using a lysate-based *in vitro* translation (PfIVT) assay where luminescence from translation of a luciferase reporter transcript provides the readout (20-22), and DDD107498 translation inhibition IC_50s_ was 60.5 nm in the 96wp assay (22) and 100 nm in the 384wp assay (20). In contrast, the DDD107498 antiplasmodial EC_50_ against *P. falciparum* ABS in standard 48h re-invasion assays is around 1 nM (Baragana 2015, Tamaki 2022). Tamaki et al suggest that this disconnect, seen with several of their PfIVT hits as well as known other known translation inhibitors, is most likely due to the assay conditions required for the PfIVT assay masking the true translation inhibition potency of some compounds (20). Our results using *in cellulo* assays, where *P. berghei* translation of native transcripts is quantified in the normal cellular context suggest this is not the complete story. DDD107498 antiplasmodial EC_50_ in *P. berghei* LS assays, where biomass reduction is quantified via a luciferase reporter after 46-48 hpi of compound treatment, is in the 1-2 nM range (16, 35, 36), while the translation inhibition EC_50_s in our cellular assay were 11.2, 13.4, and 14.5 nM (23, 35). MMV667494 sits in the middle, with an essentially identical antiplasmodial and translation inhibition efficacy (Figs. 4D-E). While the evidence that *P. falciparum* ABS resistance to DDD107498 is mediated by mutations in PfeEF2 is extremely strong (16, 37, 38), inhibition of protein synthesis *per se* may not be the ultimate, as opposed to proximate, cause of its antiplasmodial activity. Additional investigation into the relationship between translation inhibition and antiplasmodial activity within the quinoline-4-carboxamide series may help to shed light on how these compounds ultimately exert antiplasmodial activity.

The PfIVT screen of the Pathogen Box identified 19 compounds as primary hits, exerting at least 50% inhibition at a 30μM test concentration; two of these were found to be false positives based on a luciferase reporter counterscreen, while six were considered to be viable hits based on based selective inhibition of *P. falciparum* vs. human protein synthesis after a second counterscreen using *H. sapiens* lysate translation of the same luciferase reporter (20). Two of these six viable hits were MMV634140 and MMV667494, the DDD107498 analogues that were the sole hits in our *in cellulo* screen using a 5 μM screening concentration. The other four viable hits from the PfIVT Pathogen Box screen were clearly inactive in our screen (Table S2). Of these four, MMV019189, identified in the large GSK ABS antiplasmodial activity screen as TCMDC-123736 (39), which has antiplasmodial activity against both *P. falciparum* asexual blood stage parasites and *P. berghei* liver stages with EC_50s_ in the range of 750-1100 nM (33, 40), was selected for follow up interrogation at a higher concentration. Based on the roughly 10-fold difference we have seen with DDD107498 antiplasmodial vs. translation inhibition potency, MMV019189 activity against liver stage protein synthesis might have been missed with our 5μM screening concentration, as the OPP HCI assay was not intended to detect compounds with weak activity at the screening concentration. MMV019189 was tested at 25μM in our standard 30-minute coOPP assay with ACFM imaging, and inhibited both liver stage translation and HepG2 protein synthesis by a modest ∼20% (Fig. S3). With these results, it seems clear that any antiplasmodial effects MMV019189 exerts on *P. berghei* liver stages are not mediated via inhibition of parasite translation, and makes it less likely that its antiplasmodial activity against *P. falciparum* ABSs is due to inhibition of protein synthesis. The idea of targeting core cellular processes like translation is rooted in their demonstrated requirement across different *Plasmodium* species and stages, such that these compounds should be multistage active antiplasmodials. While we have shown that the threonyl tRNA synthetase inhibitor borrelidin (41, 42) has limited effects on *P. berghei* liver stage translation, and only at concentrations where the compound was cytotoxic to the HepG2 cells (23), it is the single example of low activity or inactivity seen to date amongst the validated *Plasmodium*-specific or pan-eukaryotic translation inhibitors we have tested. MMV019266, a thienopyridimide thought to target the cytoplasmic isoleucyl tRNA synthetase (43) was not identified as active in a PfIVT screen of the Malaria Box (21), but we clearly identify it as an inhibitor of native liver stage translation using the OPP assay in either HCI (Fig. 2C) or ACFM (23) modalities. The OPP HCI assay’s 0.5% hit rate for the Pathogen Box, a compound collection consisting entirely of known bioactive compounds targeting a range of pathogens, demonstrates the striking specificity of the assay, which is all the more remarkable given the low magnification, low resolution, pseudo-widefield imaging strategy we used. The OPP HCI assay was designed to provide a medium throughput, secondary screen to identify translation inhibitors amongst the many antiplasmodial hits identified from high throughput screens, or for primary screens of small, high-value compound libraries. While interrogation of antiplasmodial compound mechanisms has been almost exclusively limited to the blood stages, liver stage testing might be ideal for identifying multistage and multi-species active antiplasmodials that inhibit native cytoplasmic protein synthesis via the OPP HCI assay.

### Conclusions

Taken together, our screening results demonstrate the clear utility and reliability of the OPP HCI assay for identifying compounds that inhibit native *Plasmodium* liver stage translation and determining their selectivity in a single well. The quinoline-4-carboxamide chemotype, of which DDD107498/cabamiquine is an exemplar, deserves deeper interrogation to understand how these compounds, with widely varying relationships between their liver stage antiplasmodial efficacy and translation inhibition efficacy, ultimately kill *Plasmodium* parasites.

## Materials and Methods

### HepG2 cell seeding and direct infection with *P. berghei* sporozoites

HepG2 cells were cultured as described in detail in (26). Prior to HepG2 cell seeding, the wells of a 384wp (Falcon, cat: 353963) were filled with 20 μL of infection medium (iDMEM) comprised of DMEM (Gibco, cat: 10313-021) media supplemented with 10% FBS (v/v)(Gibco, cat: 166140-071), 1% GlutaMax (v/v)(Gibco, cat: 35050-061), 1% penicillin streptomycin (v/v)(Gibco, cat: 15140-122), 1% kanamycin sulfate (v/v)(Corning, cat: 30-006-CF), 1% penicillin-streptomycin-neomycin (v/v)(Gibco, cat: 15640-055), 0.3% Amphotericin B (v/v)(Gibco, cat: 15290-026), 0.1% gentamycin (v/v)(Gibco, cat: 15750-060). HepG2 human hepatoma cells were trypsinized, quantified with a Neubauer counting chamber, and re-suspended in iDMEM at a concentration of 175 cells/μL. 40 μL of this cell suspension was added to each of the media containing wells of the 384 wp to achieve a final concentration of 7,000 HepG2 cells per well and incubated overnight at 37 °C at 5% CO_2_. Prior to dissection, *Anopheles stephensii* mosquitos were cold anesthetized, dipped into 70% ethanol and washed twice in 1x HBSS (Corning, cat: 21-022-CV); isolated salivary glands were transferred to unsupplemented DMEM kept on ice. Sporozoites (*P. berghei* ANKA, strain 676m1c11 (44)) were released from salivary glands using a sterile pestle and resuspended before being passed through a 40 μm filter to remove debris. Sporozoites were quantified with a Neubauer counting chamber and resuspended in iDMEM at a concentration of 250 sporozoites/μL. 20 μL of the sporozoite solution was added to each well of the 384wp to achieve a final concentration of 5,000 sporozoites/well and the plates were centrifuged at 3000 rpm (2000 x *g*) for 5 minutes and returned to the incubator conditions previously described for 2 hours. At 2 hours post infection (hpi), 50 μL of infection media was removed from each well and replaced with 40 μL of fresh infection media; plates were returned to the incubator until treatment.

### CoOPP treatment and fixation

Pre-spotted compound plates were thawed at room temperature for 30 minutes, then iDMEM was dispensed to each well to achieve a 16X (80 μM) concentration, and was mixed by automated pipettor after 15 minutes. 30 μL from each well was transferred to a round-bottom 96wp (Corning, cat: 3799) and combined with 30 μL of 16X (320 μM) o-propargyl puromycin (OPP) (Invitrogen, cat: 10459) in iDMEM media to achieve an 8x working concentration for compound and OPP. The anisomycin and DDD107498 controls were not present on the original compound library plates, so were prepared separately in 1.6 mL microcentrifuge tubes, and transferred to the coOPP treatment plates with the same volumes/concentrations described for the screening compounds. For controls, each plate contained sixteen DMSO wells, eight wells of DDD107498 (1μM), and eight wells of anisomycin (10 μM). The coOPP treatment plates were incubated for 30 minutes in the 37 °C, 5% CO_2_ incubator. At 27.5 hpi, assay plates containing the infected monolayers were treated by dispensing 10 μL of pre-warmed coOPP treatments to each well, then the plates were returned to the incubator for 30 minutes. Following treatment, plates were removed from the incubator and fixed with direct addition of 26.6 μL of 16% PFA (final concentration of 4% PFA) to each well at room temperature for 15 minutes. After fixation, PFA-containing media was removed and the plates were washed three times with 100 μL of 1x PBS.

### Click chemistry and immunostaining

Samples were permeabilized with 0.5% Triton X100 (Sigma, cat: D8537) in PBS (v/v) at room temperature for 20 minutes, washed twice with PBS and blocked for ∼10 minutes in 2% BSA (Fisher, cat: 9048-46-8) in PBS (w/v) while the click chemistry (CC) labeling reaction (Vector labs, cat: CCT-1317) was prepared, according to manufacturer’s recommendation. Fluorophore addition was performed using freshly prepared CC solution for 30 minutes at room temperature, shielded from light. After the click reaction, samples were washed once with CC wash buffer and once with PBS prior to 1 h incubation in blocking solution at RT. Primary antibody labeling of *P. berghei* EEFs was performed using mouse monoclonal α-PbHSP70 (45) in 2% blocking solution (1:200) at 4 °C overnight, followed by 1 h secondary labeling with donkey α-mouse Alexafluor488 (1:500)(Invitrogen, cat: A21202) at RT, with Hoechst 33342 (1:1000) (Thermo Scientific, cat: 62249) used for labeling DNA.

### Image acquisition, segmentation and feature extraction

For the OPP HCI, pseudo-widefield image sets were acquired on a Leica SP8 confocal microscope equipped with MatrixScreener using a 10x air objective and pinhole size of 283.1 μm (approximately 4 AU) with 2x digital zoom; identical acquisition settings were used for all samples. Sequential 1024x1024 pixel images were acquired of the EEF marker, DNA, and OPP-labeled nascent proteome channels, with 4 non-overlapping fields acquired per well. ACFM images (Figs. 3C, 4C) were acquired sequentially for each channel with a 63X, 1.2 NA water objective.

CellProfiler (V2.1.1 6c2d896) was used for image segmentation and feature extraction, using identical module settings between samples. To separate the nascent proteome signal of the EEFs from the HepG2, EEFs were identified by global thresholding of the αPbHSP70 image and defined as EEF objects. These EEF objects were used to mask the OPP-A555 image to extract fluorescence intensity of the nascent proteome in HepG2 and an inverse mask was used to extract fluorescence intensity of parasite nascent proteomes. To validate that the OPP-A555 signal measured in HCI images was not influenced by signal associated to the HepG2, EEF objects were shrunken by varying degrees (Figure 2A) prior to OPP-A555 image masking and the translation intensity correlation was compared to the same samples imaged by ACFM and processed as previously described (35). The EEF objects that were shrunk down to a single, central pixel provided the highest correlation to ACFM data set, and this strategy was exclusively used in the 384wp pilot screens. All OPP mean fluorescence intensities (MFI) were normalized to the mean of the in-plate DMSO controls.

### Live luciferase assay

384 well plates were seeded with 7000 HepG2 cells per well and were directly infected with 3000 sporozoites per well as described above. After the samples were centrifuged and placed in the incubator, both compound hits from the Pathogen Box compound set were prepared in 5-point, 5-fold concentration response at 4x working concentration starting at 4 μM. At 2 hpi, the infection media in each well was reduced to 30 μL and replaced with 30 μL of fresh infection media and 20 μL of the 4x compound treatments and DMSO controls. Equal concentrations of DMSO (1:1000 v/v) were maintained across the plate. At 48 hpi, treatment media was removed by centrifugation and replaced with 50 μL of D-luciferin infection media (150 μg/mL) as in (32) and returned to the incubator for 10 minutes. After incubation, luminescence was read on a BMG Labtech Clariostar Plus plate reader and the luminescence values were normalized to the mean value of the DMSO control wells.

### Statistics and data analysis

Concentration response curve fits and EC_50_ values were generated in Prism (v7.0d) using a four parameter non-linear regression model as previously described (23), with the top of the curve fixed at 100 and the bottom of the curve fit open, except for the live luciferase assay in Fig. 4D, where the maximal effect was constrained to zero. All other statistical analysis and data handling was performed using KNIME (v4) (46). Scatter plots and box plots were generated using Plotly Chart Studio.

## Acknowledgements

We thank the University of Georgia SporoCore for providing *P. berghei-*infected mosquitos, and Medicines for Malaria Venture for providing the MMV Validation Set, Pathogen Box, and MMV634140 and MMV667494. This work was funded by NIH R21AI149275 to KKH. JLM received financial support from a South Texas Center for Emerging Infectious Diseases fellowship.

## References

1. Prudêncio M, Rodriguez A, Mota MM. 2006. The silent path to thousands of merozoites: the Plasmodium liver stage. Nature Reviews Microbiology 4:849–856.

2. Baer K, Klotz C, Kappe SH, Schnieder T, Frevert U. 2007. Release of hepatic Plasmodium yoelii merozoites into the pulmonary microvasculature. PLoS Pathog 3:e171.

3. Sturm A, Amino R, van de Sand C, Regen T, Retzlaff S, Rennenberg A, Krueger A, Pollok JM, Menard R, Heussler VT. 2006. Manipulation of host hepatocytes by the malaria parasite for delivery into liver sinusoids. Science 313:1287–90.

4. Graewe S, Rankin KE, Lehmann C, Deschermeier C, Hecht L, Froehlke U, Stanway RR, Heussler V. 2011. Hostile takeover by Plasmodium: reorganization of parasite and host cell membranes during liver stage egress. PLoS Pathog 7:e1002224.

5. Haldar K, Murphy SC, Milner DA, Taylor TE. 2007. Malaria: mechanisms of erythrocytic infection and pathological correlates of severe disease. Annu Rev Pathol 2:217–49.

6. Meibalan E, Marti M. 2017. Biology of Malaria Transmission. Cold Spring Harbor perspectives in medicine 7:a025452.

7. Burrows JN, Burlot E, Campo B, Cherbuin S, Jeanneret S, Leroy D, Spangenberg T, Waterson D, Wells TN, Willis P. 2014. Antimalarial drug discovery – the path towards eradication. Parasitology 141:128–139.

8. Burrows JN, Duparc S, Gutteridge WE, Hooft van Huijsduijnen R, Kaszubska W, Macintyre F, Mazzuri S, Möhrle JJ, Wells TNC. 2017. New developments in anti-malarial target candidate and product profiles. Malaria Journal 16:26.

9. Hovlid ML, Winzeler EA. 2016. Phenotypic Screens in Antimalarial Drug Discovery. Trends Parasitol 32:697–707.

10. Goodman CD, Su V, McFadden GI. 2007. The effects of anti-bacterials on the malaria parasite. Molecular and Biochemical Parasitology 152:181–191.

11. Dahl EL, Rosenthal PJ. 2007. Multiple antibiotics exert delayed effects against the apicoplast. Antimicrobial Agents and Chemotherapy 51:3485–3490.

12. Jackson KE, Habib S, Frugier M, Hoen R, Khan S, Pham JS, Ribas de Pouplana L, Royo M, Santos MA, Sharma A, Ralph SA. 2011. Protein translation in Plasmodium parasites. Trends Parasitol 27:467–76.

13. Jackson KE, Pham JS, Kwek M, De Silva NS, Allen SM, Goodman CD, McFadden GI, Ribas de Pouplana L, Ralph SA. 2012. Dual targeting of aminoacyl-tRNA synthetases to the apicoplast and cytosol in Plasmodium falciparum. International Journal for Parasitology 42:177–186.

14. Xie S, Griffin MDW, Winzeler EA, Ribas de Pouplana L, Tilley L. 2023. Targeting Aminoacyl tRNA Synthetases for Antimalarial Drug Development. Annu Rev Microbiol doi:10.1146/annurev-micro-032421-121210.

15. Pham JS, Dawson KL, Jackson KE, Lim EE, Pasaje CF, Turner KE, Ralph SA. 2014. Aminoacyl-tRNA synthetases as drug targets in eukaryotic parasites. Int J Parasitol Drugs Drug Resist 4:1–13.

16. Baragana B, Hallyburton I, Lee MC, Norcross NR, Grimaldi R, Otto TD, Proto WR, Blagborough AM, Meister S, Wirjanata G, Ruecker A, Upton LM, Abraham TS, Almeida MJ, Pradhan A, Porzelle A, Luksch T, Martinez MS, Luksch T, Bolscher JM, Woodland A, Norval S, Zuccotto F, Thomas J, Simeons F, Stojanovski L, Osuna-Cabello M, Brock PM, Churcher TS, Sala KA, Zakutansky SE, Jimenez-Diaz MB, Sanz LM, Riley J, Basak R, Campbell M, Avery VM, Sauerwein RW, Dechering KJ, Noviyanti R, Campo B, Frearson JA, Angulo-Barturen I, Ferrer-Bazaga S, Gamo FJ, Wyatt PG, Leroy D, Siegl P, Delves MJ, Kyle DE, et al. 2015. A novel multiple-stage antimalarial agent that inhibits protein synthesis. Nature 522:315–20.

17. McCarthy JS, Yalkinoglu Ö, Odedra A, Webster R, Oeuvray C, Tappert A, Bezuidenhout D, Giddins MJ, Dhingra SK, Fidock DA, Marquart L, Webb L, Yin X, Khandelwal A, Bagchus WM. 2021. Safety, pharmacokinetics, and antimalarial activity of the novel plasmodium eukaryotic translation elongation factor 2 inhibitor M5717: a first-in-human, randomised, placebo-controlled, double-blind, single ascending dose study and volunteer infection study. The Lancet Infectious Diseases 21:1713–1724.

18. van der Plas JL, Kuiper VP, Bagchus WM, Bodding M, Yalkinoglu O, Tappert A, Seitzinger A, Spangenberg T, Bezuidenhout D, Wilkins J, Oeuvray C, Dhingra SK, Thathy V, Fidock DA, Smidt LCA, Roozen GVT, Koopman JPR, Lamers OAC, Sijtsma J, van Schuijlenburg R, Wessels E, Meij P, Kamerling IMC, Roestenberg M, Khandelwal A. 2023. Causal chemoprophylactic activity of cabamiquine against Plasmodium falciparum in a controlled human malaria infection: a randomised, double-blind, placebo-controlled study in the Netherlands. Lancet Infect Dis 23:1164–1174.

19. Xie SC, Metcalfe RD, Dunn E, Morton CJ, Huang SC, Puhalovich T, Du Y, Wittlin S, Nie S, Luth MR, Ma L, Kim MS, Pasaje CFA, Kumpornsin K, Giannangelo C, Houghton FJ, Churchyard A, Famodimu MT, Barry DC, Gillett DL, Dey S, Kosasih CC, Newman W, Niles JC, Lee MCS, Baum J, Ottilie S, Winzeler EA, Creek DJ, Williamson N, Parker MW, Brand S, Langston SP, Dick LR, Griffin MDW, Gould AE, Tilley L. 2022. Reaction hijacking of tyrosine tRNA synthetase as a new whole-of-life-cycle antimalarial strategy. Science 376:1074–1079.

20. Tamaki F, Fisher F, Milne R, Terán FS-R, Wiedemar N, Wrobel K, Edwards D, Baumann H, Gilbert IH, Baragana B, Baum J, Wyllie S. 2022. High-Throughput Screening Platform To Identify Inhibitors of Protein Synthesis with Potential for the Treatment of Malaria. Antimicrobial Agents and Chemotherapy 66:e00237–22.

21. Ahyong V, Sheridan CM, Leon KE, Witchley JN, Diep J, DeRisi JL. 2016. Identification of Plasmodium falciparum specific translation inhibitors from the MMV Malaria Box using a high throughput in vitro translation screen. Malar J 15:173.

22. Sheridan CM, Garcia VE, Ahyong V, DeRisi JL. 2018. The Plasmodium falciparum cytoplasmic translation apparatus: a promising therapeutic target not yet exploited by clinically approved anti-malarials. Malar J 17:465.

23. McLellan JL, Sausman W, Reers AB, Bunnik EM, Hanson KK. 2023. Single-cell quantitative bioimaging of P. berghei liver stage translation. mSphere doi:10.1128/msphere.00544-23:e0054423.

24. Liu J, Xu Y, Stoleru D, Salic A. 2012. Imaging protein synthesis in cells and tissues with an alkyne analog of puromycin. Proc Natl Acad Sci U S A 109:413–8.

25. Tischer C, Hilsenstein V, Hanson K, Pepperkok R. 2014. Chapter 26 - Adaptive fluorescence microscopy by online feedback image analysis, p 489-503. In Waters JC, Wittman T (ed), Methods in Cell Biology, vol 123. Academic Press.

26. McLellan JL, Garcia-Vilanova A, Hanson KK. 2024. An Optimized P. berghei Liver Stage– HepG2 Infection Model for Simultaneous Quantitative Bioimaging of Host and Parasite Nascent Proteomes. Bio-protocol 14:e4952.

27. Schneider-Poetsch T, Ju J, Eyler DE, Dang Y, Bhat S, Merrick WC, Green R, Shen B, Liu JO. 2010. Inhibition of eukaryotic translation elongation by cycloheximide and lactimidomycin. Nat Chem Biol 6:209–217.

28. Garreau de Loubresse N, Prokhorova I, Holtkamp W, Rodnina MV, Yusupova G, Yusupov M. 2014. Structural basis for the inhibition of the eukaryotic ribosome. Nature 513:517–522.

29. Plouffe D, Brinker A, McNamara C, Henson K, Kato N, Kuhen K, Nagle A, Adrián F, Matzen JT, Anderson P, Nam TG, Gray NS, Chatterjee A, Janes J, Yan SF, Trager R, Caldwell JS, Schultz PG, Zhou Y, Winzeler EA. 2008. In silico activity profiling reveals the mechanism of action of antimalarials discovered in a high-throughput screen. Proc Natl Acad Sci U S A 105:9059–64.

30. Meister S, Plouffe DM, Kuhen KL, Bonamy GMC, Wu T, Barnes SW, Bopp SE, Borboa R, Bright AT, Che JW, Cohen S, Dharia NV, Gagaring K, Gettayacamin M, Gordon P, Groessl T, Kato N, Lee MCS, McNamara CW, Fidock DA, Nagle A, Nam TG, Richmond W, Roland J, Rottmann M, Zhou B, Froissard P, Glynne RJ, Mazier D, Sattabongkot J, Schultz PG, Tuntland T, Walker JR, Zhou YY, Chatterjee A, Diagana TT, Winzeler EA. 2011. Imaging of Plasmodium Liver Stages to Drive Next-Generation Antimalarial Drug Discovery. Science 334:1372–1377.

31. Hoepfner D, McNamara CW, Lim CS, Studer C, Riedl R, Aust T, McCormack SL, Plouffe DM, Meister S, Schuierer S, Plikat U, Hartmann N, Staedtler F, Cotesta S, Schmitt EK, Petersen F, Supek F, Glynne RJ, Tallarico JA, Porter JA, Fishman MC, Bodenreider C, Diagana TT, Movva NR, Winzeler EA. 2012. Selective and specific inhibition of the plasmodium falciparum lysyl-tRNA synthetase by the fungal secondary metabolite cladosporin. Cell Host Microbe 11:654–63.

32. Van Voorhis WC, Adams JH, Adelfio R, Ahyong V, Akabas MH, Alano P, Alday A, Alemán Resto Y, Alsibaee A, Alzualde A, Andrews KT, Avery SV, Avery VM, Ayong L, Baker M, Baker S, Ben Mamoun C, Bhatia S, Bickle Q, Bounaadja L, Bowling T, Bosch J, Boucher LE, Boyom FF, Brea J, Brennan M, Burton A, Caffrey CR, Camarda G, Carrasquilla M, Carter D, Belen Cassera M, Chih-Chien Cheng K, Chindaudomsate W, Chubb A, Colon BL, Colón-López DD, Corbett Y, Crowther GJ, Cowan N, D’Alessandro S, Le Dang N, Delves M, DeRisi JL, D. Ay, Duffy S, Abd El-Salam El-Sayed S, Ferdig MT, Fernández Robledo JA, Fidock DA, et al. 2015. Open Source Drug Discovery with the Malaria Box Compound Collection for Neglected Diseases and Beyond. PLOS Pathogens 12:e1005763.

33. MMV. Pathogen Box Platemaps with Biological Activity Data. https://www.mmv.org/sites/default/files/uploads/docs/mmv_open/Pathogen_Box_Activity_Biological_Data_Smiles.xlsx. Accessed May 25, 2024.

34. Baragaña B, Norcross NR, Wilson C, Porzelle A, Hallyburton I, Grimaldi R, Osuna-Cabello M, Norval S, Riley J, Stojanovski L, Simeons FRC, Wyatt PG, Delves MJ, Meister S, Duffy S, Avery VM, Winzeler EA, Sinden RE, Wittlin S, Frearson JA, Gray DW, Fairlamb AH, Waterson D, Campbell SF, Willis P, Read KD, Gilbert IH. 2016. Discovery of a Quinoline-4-carboxamide Derivative with a Novel Mechanism of Action, Multistage Antimalarial Activity, and Potent in Vivo Efficacy. Journal of Medicinal Chemistry 59:9672–9685.

35. McLellan JL, Hanson KK. 2024. Differential effects of translation inhibitors on liver stage parasites. Life Science Alliance 7.

36. Arez F, Rebelo SP, Fontinha D, Simao D, Martins TR, Machado M, Fischli C, Oeuvray C, Badolo L, Carrondo MJT, Rottmann M, Spangenberg T, Brito C, Greco B, Prudêncio M, Alves PM. 2019. Flexible 3D Cell-Based Platforms for the Discovery and Profiling of Novel Drugs Targeting Hepatic Infection. Acs Infectious Diseases 5:1831–1842.

37. Rottmann M, Jonat B, Gumpp C, Dhingra SK, Giddins MJ, Yin X, Badolo L, Greco B, Fidock DA, Oeuvray C, Spangenberg T. 2020. Preclinical Antimalarial Combination Study of M5717, a Plasmodium falciparum Elongation Factor 2 Inhibitor, and Pyronaridine, a Hemozoin Formation Inhibitor. Antimicrob Agents Chemother 64:10.1128/aac.02181-19.

38. Stadler E, Maiga M, Friedrich L, Thathy V, Demarta-Gatsi C, Dara A, Sogore F, Striepen J, Oeuvray C, Djimde AA, Lee MCS, Dembele L, Fidock DA, Khoury DS, Spangenberg T. 2023. Propensity of selecting mutant parasites for the antimalarial drug cabamiquine. Nat Commun 14:5205.

39. Gamo FJ, Sanz LM, Vidal J, de Cozar C, Alvarez E, Lavandera JL, Vanderwall DE, Green DVS, Kumar V, Hasan S, Brown JR, Peishoff CE, Cardon LR, Garcia-Bustos JF. 2010. Thousands of chemical starting points for antimalarial lead identification. Nature 465:305–U56.

40. Duffy S, Sykes ML, Jones AJ, Shelper TB, Simpson M, Lang R, Poulsen SA, Sleebs BE, Avery VM. 2017. Screening the Medicines for Malaria Venture Pathogen Box across Multiple Pathogens Reclassifies Starting Points for Open-Source Drug Discovery. Antimicrobial Agents and Chemotherapy 61.

41. Novoa EM, Camacho N, Tor A, Wilkinson B, Moss S, Marin-Garcia P, Azcarate IG, Bautista JM, Mirando AC, Francklyn CS, Varon S, Royo M, Cortes A, Ribas de Pouplana L. 2014. Analogs of natural aminoacyl-tRNA synthetase inhibitors clear malaria in vivo. Proc Natl Acad Sci U S A 111:E5508–17.

42. Fang P, Yu X, Jeong SJ, Mirando A, Chen K, Chen X, Kim S, Francklyn CS, Guo M. 2015. Structural basis for full-spectrum inhibition of translational functions on a tRNA synthetase. Nat Commun 6:6402.

43. Istvan ES, Guerra F, Abraham M, Huang KS, Rocamora F, Zhao H, Xu L, Pasaje C, Kumpornsin K, Luth MR, Cui H, Yang T, Palomo Diaz S, Gomez-Lorenzo MG, Qahash T, Mittal N, Ottilie S, Niles J, Lee MCS, Llinas M, Kato N, Okombo J, Fidock DA, Schimmel P, Gamo FJ, Goldberg DE, Winzeler EA. 2023. Cytoplasmic isoleucyl tRNA synthetase as an attractive multistage antimalarial drug target. Sci Transl Med 15:eadc9249.

44. Franke-Fayard B, Janse CJ, Cunha-Rodrigues M, Ramesar J, Büscher P, Que I, Löwik C, Voshol PJ, den Boer MAM, van Duinen SG, Febbraio M, Mota MM, Waters AP. 2005. Murine malaria parasite sequestration: CD36 is the major receptor, but cerebral pathology is unlinked to sequestration. Proceedings of the National Academy of Sciences of the United States of America 102:11468–11473.

45. Tsuji M, Mattei D, Nussenzweig RS, Eichinger D, Zavala F. 1994. Demonstration of heat-shock protein 70 in the sporozoite stage of malaria parasites. Parasitol Res 80:16–21.

46. Berthold MR, Cebron N, Dill F, Gabriel TR, Kötter T, Meinl T, Ohl P, Sieb C, Thiel K, Wiswedel B. KNIME: The Konstanz Information Miner, p 319-326. In (ed), Springer Berlin Heidelberg,

