## Supplementary Figures and Tables for "A high content imaging assay for identification of specific inhibitors of native *Plasmodium* liver stage protein synthesis": McLellan et al 2024 SuppFigs.pdf

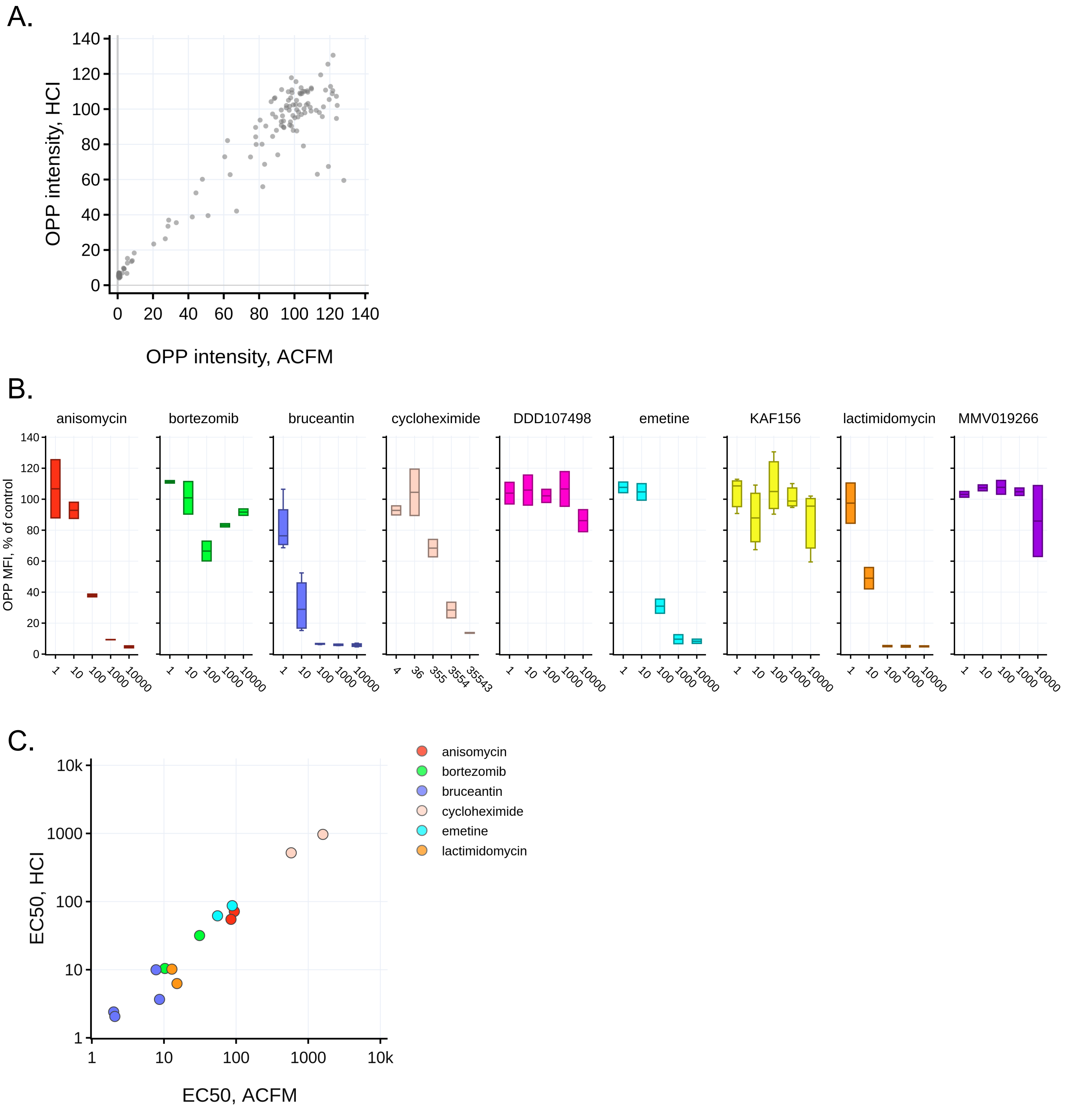


**Figure S1. Validation of OPP HCI assay for quantification of HepG2 translation. A)** HepG2 translation intensity (OPP-A555) in HCI images compared with published OPP-MFIs from ACFM images of the same wells (McLellan et al., 2023), comprising 150 wells from eight independent experiments. Each dot represents the MFI for a single well normalized to the mean of the in-plate DMSO controls, which was set to 100. **B)** Concentration response for HepG2 translation inhibition with the HCI assay (OPP MFI), normalized to DMSO controls following acute 4 h treatment with translation inhibitors labeled in each plot. **C)** Comparison of translation inhibitor potency ($EC_{50}$ value) between HCI assay and ACFM assay. For B-C, data are from 2-4 independent experiments.

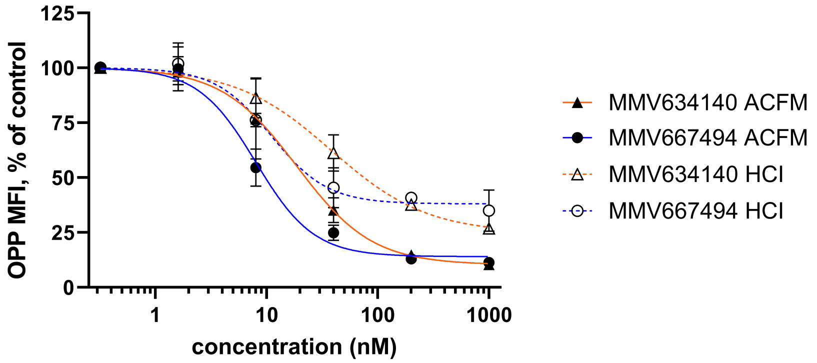


**MMV634140**
ACFM $EC_{50}$ = 18.7 nM (12.5 – 29.1 nM)
HCI $EC_{50}$ = 38.8 nM (24.0 – 74.9 nM)
ACFM % max inhibition = 89.7%
HCI % max inhibition= 75.3%

**MMV667494**
ACFM $EC_{50}$ = 8.00 nM (6.0 – 11.7 nM)
HCI $EC_{50}$ = 11.0 nM (7.3 – 18.9 nM)
ACFM % max inhibition= 86%
HCI % max inhibition= 62%

**Figure S2. ACFM validation of HCI concentration-response for Pathogen Box hits.** Concentration response curve fits and $EC_{50}$ values for *P. berghei* translation inhibition, normalized to DMSO controls following coOPP treatment with Pathogen Box compound hits from 27.5-28 hpi, comparing HCI assay data to ACFM assay data from one of the three independent experiments reported in Fig. 4D. Points  show the mean of 2 replicate wells, and bars show standard deviation. $EC_{50}$ values and associated 95% confidence intervals (in parentheses) along with % maximum inhibition are reported for both ACFM and HCI assay data..

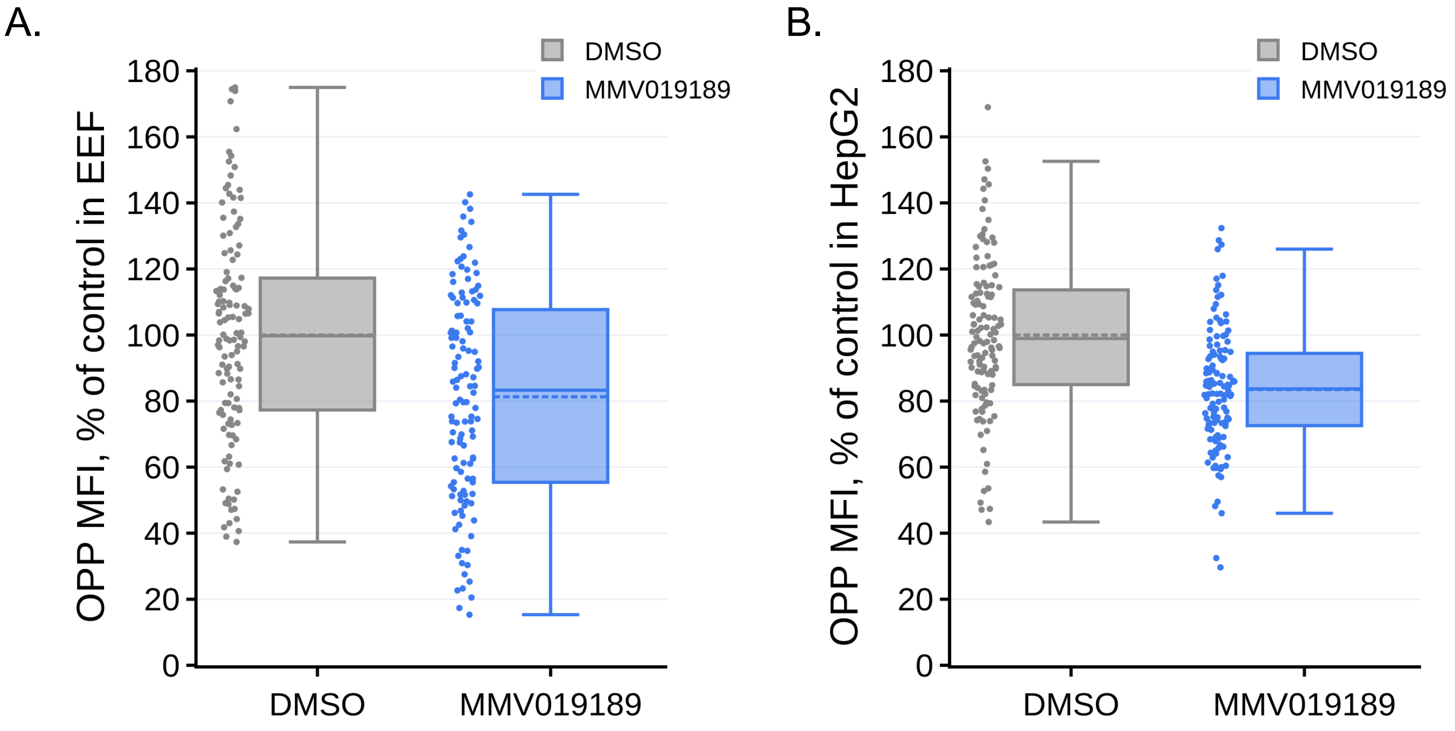


**Figure S3. Inhibition of *P. berghei* liver stage and HepG2 translation by 25 µM MMV019189. A)** *P. berghei* and **B) H**epG2 nascent proteomes (OPP MFI) following coOPP treatment with 25 µM MMV019189 from 27.5-28 hpi, quantified from single parasite ACFM images, with all data normalized to mean of the DMSO controls. Each dot represents the translational output of a single EEF (A) or that of the in-image HepG2 cells (B).
